# Epigenetic clock and DNA methylation studies of roe deer in the wild

**DOI:** 10.1101/2020.09.21.306613

**Authors:** Jean-François Lemaître, Benjamin Rey, Jean-Michel Gaillard, Corinne Régis, Emmanuelle Gilot, Maryline Pellerin, Amin Haghani, Joseph A. Zoller, Caesar Z. Li, Steve Horvath

## Abstract

DNA methylation-based biomarkers of aging (epigenetic clocks) promise to lead to new insights in the evolutionary biology of ageing. Relatively little is known about how the natural environment affects epigenetic aging effects in wild species. In this study, we took advantage of a unique long-term (>40 years) longitudinal monitoring of individual roe deer (*Capreolus capreolus*) living in two wild populations (Chizé and Trois Fontaines, France) facing different ecological contexts to investigate the relationship between chronological age and levels of DNA methylation (DNAm). We generated novel DNA methylation data from n=90 blood samples using a custom methylation array (HorvathMammalMethylChip40). We present three DNA methylation-based estimators of age (DNAm or epigenetic age), which were trained in males, females, and both sexes combined. We investigated how sex differences influenced the relationship between DNAm age and chronological age through the use of sex-specific epigenetic clocks. Our results highlight that both populations and sex influence the epigenetic age, with the bias toward a stronger male average age acceleration (i.e. differences between epigenetic age and chronological ages) particularly pronounced in the population facing harsh environmental conditions. Further, we identify the main sites of epigenetic alteration that have distinct aging patterns across the two sexes. These findings open the door to promising avenues of research at the crossroad of evolutionary biology and biogerontology.

## Introduction

The last decades have seen an increasing interest for the study of ageing in the wild (Monaghan et al. 2008; Fletcher and Selman 2015; Gaillard and Lemaître 2020). The starting point of this infatuation is undeniably the compilation of evidence reporting that - in populations of animals in the wild - senescence occurs in demographic performance (actuarial senescence: Brunet-Rossinni & Austad, 2006; Nussey et al., 2013; reproductive senescence: Lemaître & Gaillard, 2017; Nussey et al., 2013), phenotypic performance (body mass: Douhard et al., 2017; Nussey et al. 2011; foraging efficiency: Lecomte et al., 2010; MacNulty et al., 2009) and physiological traits (e.g. immune parameters, Nussey et al. 2012; haematological parameters, Jégo et al. 2014; steroid levels, Sugianto et al. 2020). Nowadays, the age-specific decline in demographic and physiological performance is considered to be the rule rather than the exception in the wild, at least in mammals and birds (Gaillard & Lemaître, 2020; Nussey et al., 2013; but see also Zajitschek et al., 2020).

Animal populations in which individuals are monitored from birth to death in the wild provide a unique (but largely untapped) resource for studying individual differences in health and mortality risk at old ages (Gaillard and Lemaître 2020; Lemaître et al. 2020b). Multiple lines of evidence emphasize the relevance of such longitudinal and individually-based data. First, the vast majority of the current research in biogerontology focused on inbred laboratory organisms with no or low genetic variation and maintained under controlled conditions (e.g. *Caenorhabditis elegans, Drosophila melanogaster*, laboratory rodents, Partridge, 2010). Studies performed on those species have led to major breakthrough in the mechanisms regulating the ageing process from the molecular to the individual level (López-Otín et al. 2013; Kennedy et al. 2014). However, their findings can be difficult to extrapolate to species living in more complex environments (Briga and Verhulst 2015), with diverse genetic background and much longer lifespan and thereby different life history strategies, such as humans (Perlman 2016). Even if studies of non-human primates kept in captive conditions are increasing (Languille et al. 2012; Jasinska 2020), the full diversity of mammalian species displaying life-history traits and life styles similar to the ones observed in humans (i.e. whether there are socially monogamous, long-lived, provide extensive periods of parental care or create tight social bounds with conspecifics) is yet to be considered. In addition, the study of the ageing process in the wild enables - by essence - to investigate the role played by the environment, an important piece of the ageing conundrum. Similar to what has been described in human populations (e.g. Robine et al., 2012 in the context of climatic variables), there is increasing evidence that environmental factors modulate ageing patterns in the wild (Nussey et al. 2007; Holand et al. 2016). For instance, it is increasingly recognized that the social environment can have a major influence on health and mortality risk at late ages (Berger et al. 2018; Snyder-Mackler et al. 2020), by notably interacting with some hallmarks of ageing (e.g. telomere dynamics in Seychelle warblers, *Acrocephalus sechellensis*, Hammers et al., 2019). In addition, while mammalian females generally live longer than males in the wild (Lemaître et al. 2020c), as commonly observed in humans or laboratory rodents (Austad and Fischer 2016; Zarulli et al. 2018), the exact mechanisms modulating these sex differences in survival are yet to be deciphered (Tower 2017; Marais et al. 2018). In that context, the focus on wild populations can be particularly relevant as the magnitude of sex differences in lifespan is likely modulated by environmental conditions, in interaction with the sex differences in genetic background (Lemaître et al. 2020c; Tidière et al. 2020). Finally, widening the scope of model species for ageing research can provide important insights for ‘healthspan extension’, notably by targeting wild animal populations displaying extended lifespan compared to the one expected for their body size (Austad 2010) and by including more appropriate senescence metrics (Lemaître et al. 2020a; Ronget and Gaillard 2020). To reach these goals, accurate markers of both chronological and biological ages on a wide range of organisms are required. The pan tissue epigenetic clock based on DNA methylation (see Horvath, 2013) is a promising indicator of biological age in humans (Paoli-Iseppi et al. 2017; Parrott and Bertucci 2019; Bell et al. 2019).

DNA methylation (DNAm) of cytosine residues within CpG dinucleotides (5-methyl-cytosine) across the genome constitutes a key epigenetic DNA modification tightly linked to the ageing process (Horvath and Raj 2018). Indeed, DNA methylation patterns accurately predict chronological age in humans (Horvath 2013; Jung and Pfeifer 2015) and captive mammals reared in laboratory conditions. Such strong relationship between age and DNAm has been found in many cell types (e.g. white blood cells, brain, liver; see Horvath, 2013). A comparative analysis of methylomes indicates that methylation can also be used to assess reliably physiological aging across mammals (Wang et al. 2020). The discrepancy between pigenetic age and chronological age (epigenetic acceleration) is associated in humans with a wide range of metabolic, infectious and degenerative diseases (Horvath et al. 2014; Horvath and Levine 2015), as well as cancer (Levine et al. 2015) and mortality (Marioni et al. 2015; Chen et al. 2016; Christiansen et al. 2016). We hypothesize that DNA methylation profiles integrates environmental effects that might modulate the pace of the epigenetic clock. To address this hypothesis. we studied epigenetic ageing in the wild and in a sex-specific way.

In this study, we took advantage of a unique long-term (>40 years) longitudinal monitoring of individual roe deer (*Capreolus capreolus*) living in two populations facing different ecological contexts to investigate the relationship between chronological age and levels of DNA methylation. All roe deer used in this study have been captured within their first year of life, when age can be accurately assigned (Hewison et al. 1999). First, we expected that the epigenetic clock built from peripheral blood leucocyte DNA should provide an accurate estimation of chronological age in roe deer in the wild. Second, we performed an epigenome wide association analysis (EWAS) to identify CpGs that were the most likely to be associated with aging in roe deer. Third, based on evidence that the pace of epigenetic age is modulated by environmental factors and provides reliable information on time to death (see above), we expected that the *Average age acceleration* (i.e. average difference between DNAm age and chronological age) would be higher in the population facing harsh environmental conditions than in the population facing favorable environmental conditions. Finally, since male roe deer show higher initial adult mortality and rate of actuarial senescence than females (Gaillard et al. 2004), we expected that the *Average age acceleration* would be higher for males than for females. Moreover, thanks to the epigenome wide association analysis, we expected to identify specific CpGs displaying sex-specific DNAm aging profiles.

## Methods

### Study populations

We sampled roe deer living in two enclosed forests with markedly different environmental contexts: Trois Fontaines (TF) and Chizé (CH). The Trois Fontaines forest (1,360 ha) is located in north-eastern France (48°43’N, 4°55’E) and is characterized by a continental climate, moderately severe winters and warm and rainy summers. This site has rich soils and provides high quality habitat for roe deer (Pettorelli et al. 2006). In contrast, the Chizé forest (2,614 ha) is located in western France (46°50’N, 0°25’W) and is characterized by temperate oceanic climate with Mediterranean influences. This site has a low productivity due to poor quality soils and frequent summer droughts (Pettorelli et al. 2006), and thereby provides a quite poor habitat for roe deer in most years. Individuals from these two populations have been intensively monitored using a long-term Capture-Mark-Recapture program since 1975 and 1977 (for Trois Fontaines and Chizé, respectively). In each site, 10-12 days of capture using drive-netting are organized every year between December and March (see Gaillard et al., 1993 for details on capture sessions), which allows capturing and measuring about half the population every year. Once a roe deer is captured, its sex and body mass (to the nearest 50g) are recorded and a basic clinical examination is performed. All individuals included in our analyses were of known age because they were either caught as newborn in spring (see Delorme et al. 1988 for further details) or as c.a. 8 months old during winter captures, when they still have their milk teeth (most often incisors and always premolars, Flerov 1952).

### Roe deer blood samples and dna extraction

In 2016 and 2017, we collected blood samples (up to 1mL per kg of body mass) from the jugular vein. Within 30 min of sampling, the blood was centrifuged at 3000 g for 10 min and the plasma layer was removed before washing the cells with an equivalent volume of 0.9% w/v NaCl solution. After a second centrifugation, the intermediate buffy coat layer, comprising mainly leukocytes, was collected in a 1.5-mL Eppendorf tube and immediately frozen at −80 °C in a portable freezer (Telstar SF 8025) until further use.

We extracted genomic DNA from leucocytes using the Macherey-Nagel NucleoSpin® Blood QuickPure kit. DNA purity was assessed using a Nanodrop ND-1000 spectrophotometer (Thermo Scientific, Wilmington DE, USA). For all samples, the purity absorption range was 1.7 - 2.0 for the 260/280 nm ratio and > 1.8 for the 260/230 nm ratio. We selected 96 samples by balancing the numbers of individuals among ages, and between populations and sexes. DNA concentration was determined spectrophotometrically using the Qubit assay kit. DNA samples were then diluted in ultrapure water to reach a concentration of ∼70 ng.µl^-1^ and displayed in a microplate to complete the DNA methylation protocol (see below). For 6 samples, the concentrations obtained after dilution were too low compared to the expected concentrations of 70 ng/µl and were excluded from the dataset. The 90 roe deer samples analysed in this study correspond to 79 individuals aged from 8 months to 13.5 years of age. This age range encompasses most of the roe deer lifespan as individuals older than 15 years of age are rarely observed in the wild (the oldest age ever recorded for a roe deer monitored in the wild being 17.5 years old, Gaillard et al. 1998).

### DNA Methylation data

We generated DNA methylation data using the custom Illumina chip “HorvathMammalMethylChip40”. The mammalian methylation array is attractive because it provides very high coverage (over thousand X) of highly conserved CpGs in mammals. Two thousand out of 38k probes were selected based on their utility for human biomarker studies: these CpGs, which were previously implemented in human Illumina Infinium arrays (EPIC, 450K) were selected due to their relevance for estimating age, blood cell counts, or the proportion of neurons in brain tissue. The remaining 35,988 probes were chosen to assess cytosine DNA methylation levels in mammalian species. Each probe is designed to cover a certain subset of species, such that overall all species have a high number of probes (Arneson, Ernst, and S. H., unpublished data). The particular subset of species for each probe is provided in the chip manifest file can be found at Gene Expression Omnibus (GEO) at NCBI as platform GPL28271. The SeSaMe normalization method was used to define beta values for each probe (Zhou et al. 2018).

### Statistical analyses

We first aimed to detect the function providing the best fit of the relationship linking DNAm age and chronological age. We thus compared three models corresponding to (1) an absence of relationship (constant model), (2) a constant increase of DNAm with age (linear model), and (3) a non-linear increase of DNAm with age (quadratic model, Table S1). The most parsimonious model was selected using the Akaike Information Criterion (AIC). We calculated AIC weights (AICw) to assess the relative likelihood that a given model was the best among the three fitted models (Burnham and Anderson 2002). We selected the model with the lowest AIC, but when the difference in AIC (denoted ΔAIC) between two competing models was less than two units, we retained the simplest model in accordance with parsimony rules (Burnham and Anderson 2002).

Second, we analyzed factors that could explain the between-sample variation in the average age acceleration. We computed the ‘Average age acceleration’ as the difference between DNAm age and chronological age following Horvath (2013). We then investigated whether the average age acceleration was influenced by sex, population and roe deer body mass (measured at capture) for the three main life stages in terms of survivorship in roe deer (Gaillard et al. 1993): juvenile (< 1 year of age), prime-age (1 to 8 years of age), and senescent (>8 years of age). For both prime-age and senescent life stages we ran a set of 14 models with the Average age acceleration as the dependent variable and sex, population and body mass as the independent variables (see Table S2 for a full list of models). To avoid fitting over-parameterized models, we did not include any three-way interactions. Due to the low number of individuals of 1 year of age (*N*=8), we only fitted 4 models for the juvenile life stage, the constant model (i.e. no detectable influence of any independent variable) and models including either a linear effect of body mass, sex differences, or population differences. In all cases, the best fitting model was selected using AIC (see above).

Third, we investigated further how sex influenced the relationship between DNAm age and chronological age through the use of sex-specific epigenetic clocks. For this purpose, we first built an epigenetic clock (called ‘female clock’) using data from females’ samples only, and then investigated the relationship between the female clock and both male and female chronological age. We performed a similar analysis with an epigenetic clock built using data from males’ samples only (called ‘male clock’). The 90 samples analyzed in our study correspond to 79 different individuals (i.e. 11 individuals were sampled both in 2016 and 2017). To account for this pseudo-replication problem (sensu Hurlbert 1984) we thus replicated all analyses using linear mixed-effects models, which included a random effect of individual roe deer, using the R-package *lme4* (Bates et al. 2015). For all models, results were qualitatively unchanged (see Electronic Supplementary Material, Table S3) and for the sake of simplicity, we only report results from simple linear models below.

Fourth, we performed an epigenome wide association study of chronological age in roe deer. Unfortunately, a good genome assembly is not available for roe deer. Therefore, we performed an EWAS analysis based on the related White-tailed deer, *Odocoileus virginianus*, (Ovir.te_1.0) genome assembly). In total, 32,767 probes from the HorvathMammalMethylChip40 were aligned to loci that are proximal to 6,314 genes in the Ovir.te_1.0 genome assembly. Due to the high inter-species conservation of the probes on the array, findings can probably be extrapolated to roe deer, and even humans or other mammalian species. To assess the potential mechanism, we used a multivariate regression model to identify the CpGs that have a distinct pattern of DNAm aging between the sexes. We used two multivariate models. In the first model, DNAm levels of an individual CpGs were regressed on sex-specific chronological age to identify the loci with DNAm aging that are shared between sexes (“Age” main effect), and also the basal sex difference that is independent of chronological age (“Sex” main effect). In the second model, we included an interaction term to identify the CpGs with distinct DNAm aging between males and females.

## Results

### Relationship between DNAm and chronological age

The model that best described the relationship between DNAm and chronological age was the quadratic model (Table 1a; **Figure 1a**; Table S1). This might be due to the fact that DNA methylation accumulates at a faster rate during the growth and development of juveniles than later in life when roe deer have reached their full size (at 2 years of age, roe deer have reached > 90% of their asymptotic mass, Hewison et al. 2011). Accordingly, the model that best described the relationship between DNAm and chronological age in adults only (2 to 14 years of age) was the linear model (slope of 0.79 ± 0.02, *N*=82, R^2^ = 0.93; **Figure 1b**; Table S1). For a subset of 25 adults from the Trois-Fontaines population, the exact date of birth was known meaning that it was possible to compute age at a very fine-scale resolution (i.e. in days). The use of this chronological age data measured in days slightly improved the fit of the relationship compared to the one obtained with the chronological age measured in years but the slope was left unchanged (slope of 0.79 ± 0.04, *N*= 25, R^2^ = 0.95; **Figure 1c**).

**Table 1:**
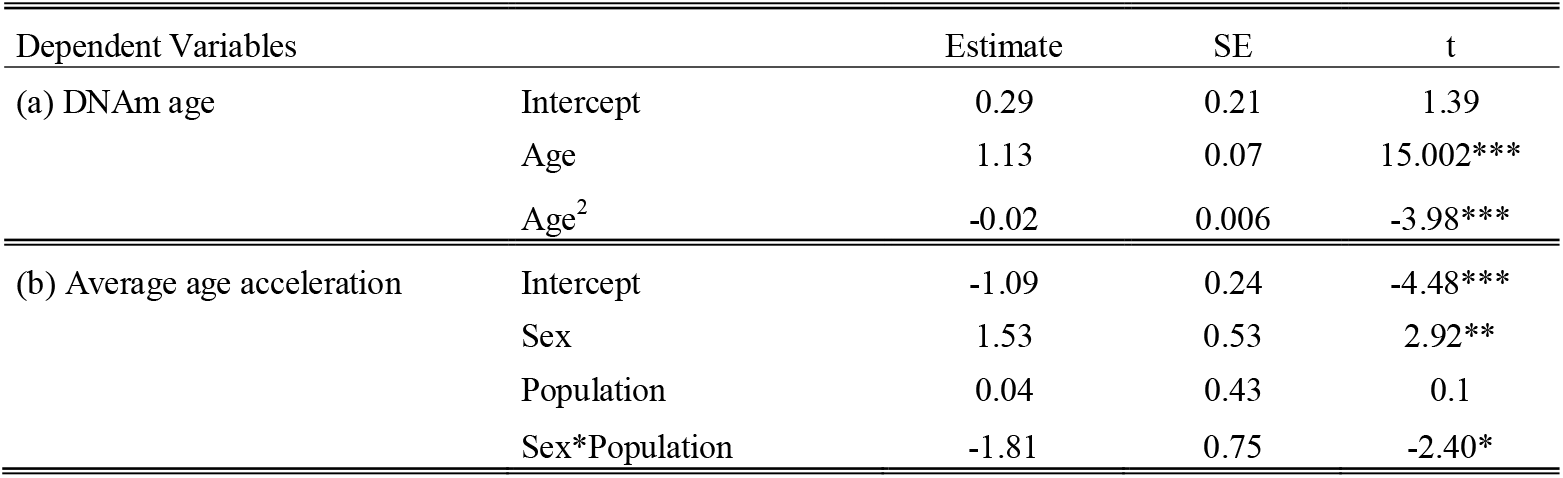
Parameters of the selected models discussed in the main text. (a) Quadratic model describing the relationship between DNAm age and chronological age in roe deer (*N*=90; R^2^ = 0.95). (b) Best model explaining variation in the Average age acceleration for senescent roe deer (i.e. >8 years of age) (*N*=23; R^2^ = 0.35) (\**p*<0.05, \*\**p*<0.01,*** *p*< 0.001).

**Figure 1:**
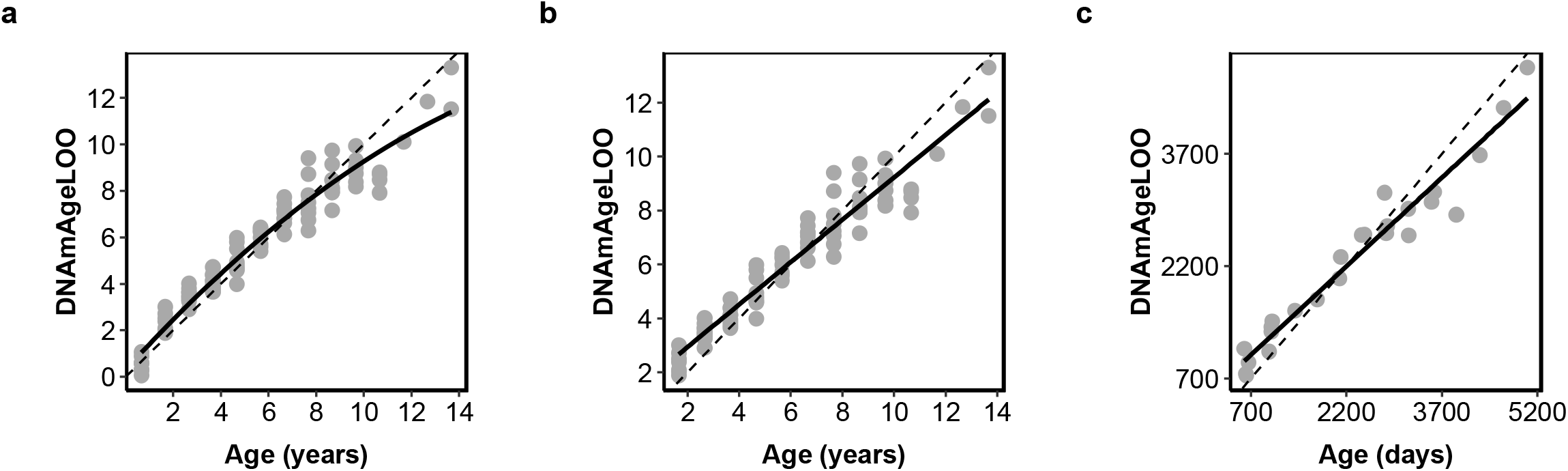
Epigenetic clock for roe deer built with individual methylation profiles from white blood cells DNA. (a) Epigenetic clock for known-aged individuals between 8 months and 14 years of age (*N*=90; R^2^=0.95); (b) Epigenetic clock for adult roe deer (i.e. >1 year old, *N*=82; R^2^=0.93); (c) Epigenetic clock for individuals where the exact age in days was known (*N*=25; R^2^=0.95). The epigenetic age (DNAmAgeLoo) is expressed in years. In all graphs, the dashed line corresponds to the regression line y = x.

### Factors affecting the average age acceleration

When focusing on juveniles only, the average age acceleration was best explained by body mass (Table S2) as shown by the positive relationship between these two variables (slope of 0.09 ± 0.03, R^2^ = 0.59; *N*=8, **Figure 2a**). On the contrary, for adults, the constant model of the average age acceleration was selected, even if we observed a trend for a higher average acceleration in males than in females (difference in intercepts of 0.36 ± 0.20, *N*=82, **Figure 2b**). We then investigated in more details the factors potentially explaining the average age acceleration in adults by running separate analyses for prime-age and senescent adults. In prime-aged adults, the selected model was again the constant model of the average age acceleration, whereas in senescent individuals, the selected model included a sex by population interaction (Table S2). The Average age acceleration was higher in males at Chizé (0.44 ± 0.37) than both in females at Chizé (−1.06 ± 0.23) and in males at Trois-Fontaines (- 1.32 ± 0.14) (Table 1b; **Figure 2c**).

**Figure 2:**
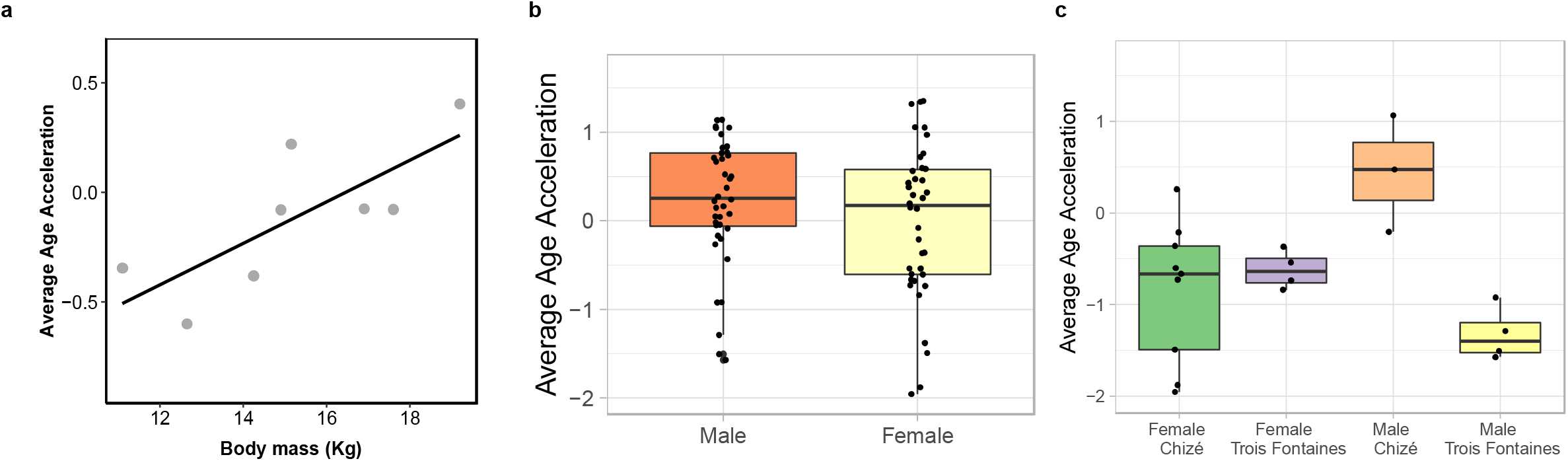
Average age acceleration in the epigenetic clock of wild roe deer. (a) Relationship between the average age acceleration and the body mass for juveniles (i.e. 8 months, *N*=8), (b) Sex-differences in the *average age acceleration* between adult males and adult females (i.e. > 1 years old, *N*=82) (c) Average age acceleration for senescent individuals (i.e. > 8 years old, *N*=23) split by sex and population.

### Influence of sex on the relationship between DNAm age and chronological age in adults

When using the female clock, a trend for interactive effects of age and sex occurred (Table 2a). As expected, the fit was much better in females (slope of 0.85 ± 0.01, *N*=42, R^2^=0.99) than in males (slope of 0.76 ± 0.05, *N*=40, R^2^=0.86). Interestingly, most male’s DNAm located above the line where DNAm age exactly matched chronological age, meaning that males are biologically older than their chronological age estimated using the female clock (**Figure 3a**). The opposite pattern occurred when using the male clock (Table 2b, **Figure 3b**). Beyond 7 years of age, females were consistently biologically younger than their chronological age estimated from the male clock (**Figure 3b**). As expected, the fit was also much better for males (slope of 0.73 ± 0.02, *N*=40, R^2^=0.98) than for females (slope of 0.48 ± 0.03, *N*=42, R^2^=0.86) when using the male clock.

**Table 2:**
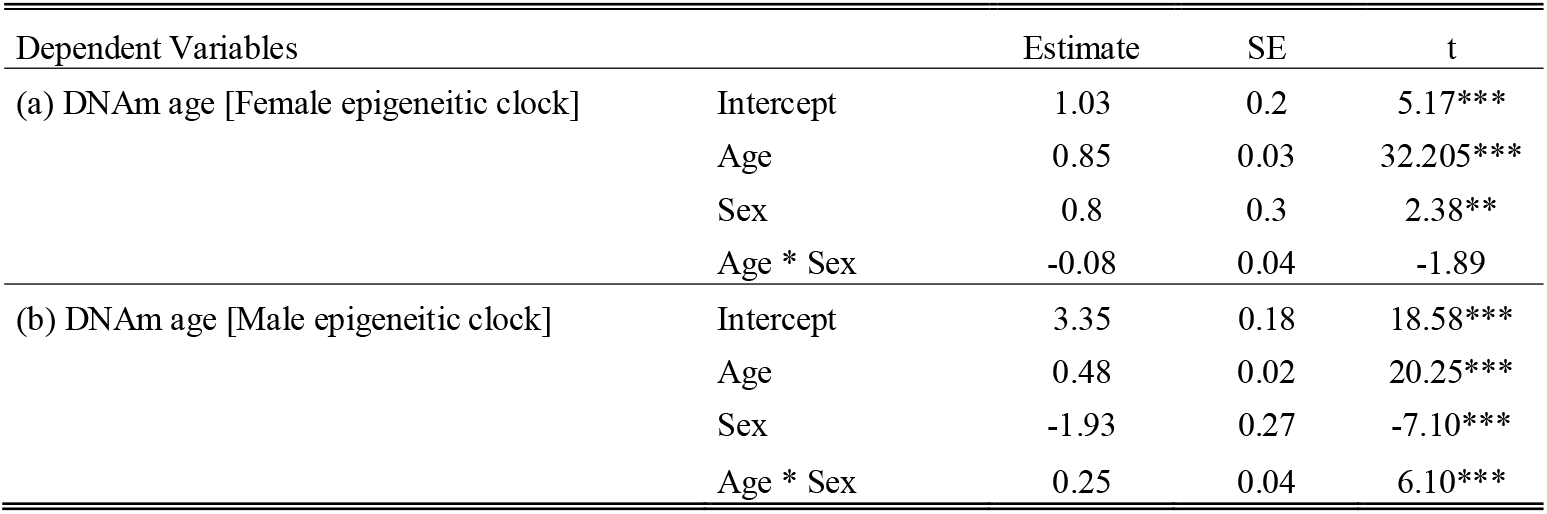
Parameters of the models testing for an interaction between chronological age and sex using the female (a) or (b) the male clock (\**p*<0.05, \*\**p*<0.01,*** *p*< 0.001).

**Figure 3:**
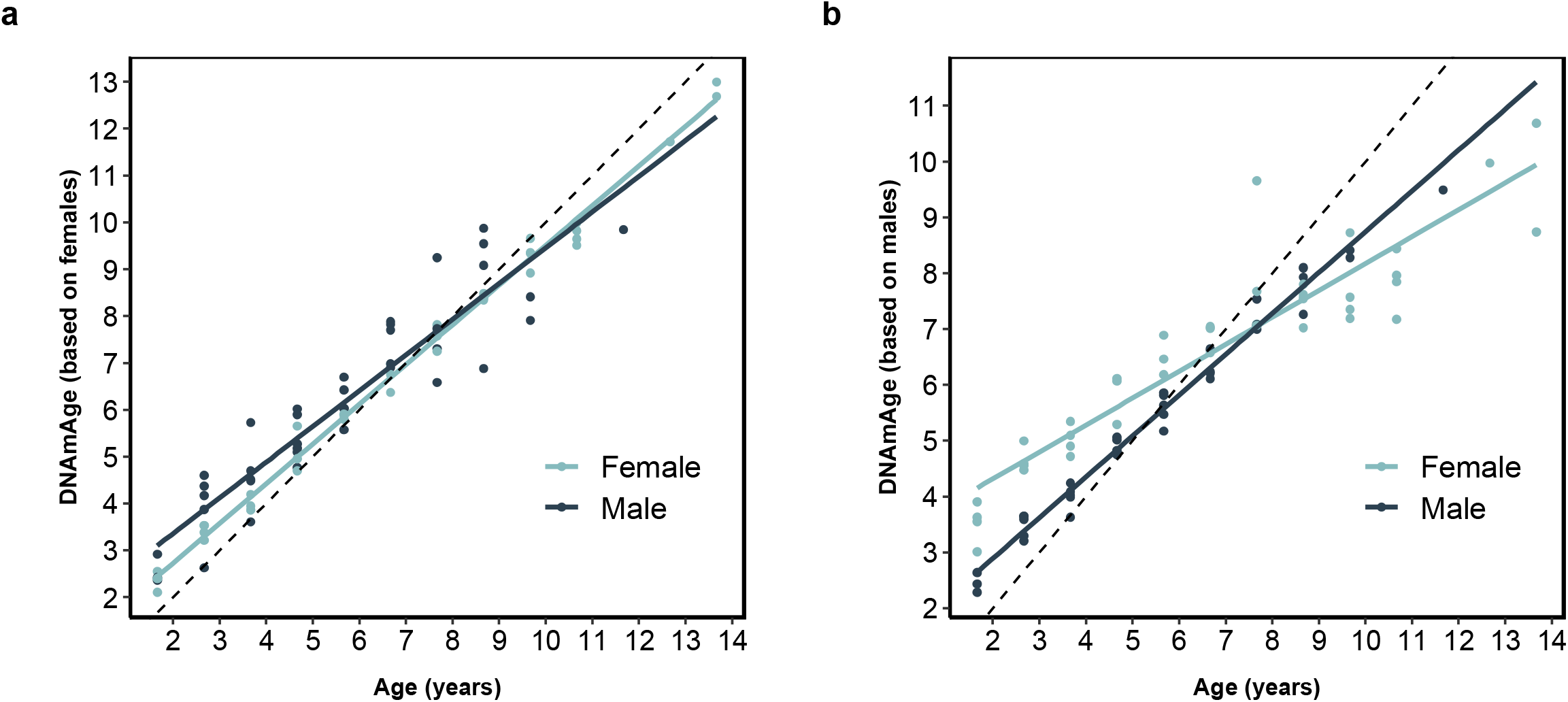
Sex-specific epigenetic clock in roe deer. (a) Relationship between DNAm age estimated with the female clock and both male and female chronological ages (b) Relationship between DNAm age estimated with the male clock and both male and female chronological ages. In all graphs, the dashed line corresponds to the regression line y = x. Females are displayed in light blue and males in dark blue.

### Epigenome wide association study of chronological age

The EWAS results revealed that chronological age alters DNAm in a large number of Loci (**Figure 4**). At a genome significance level (p<10^−8^), a total of 1992 loci showed DNAm aging. The top affected CpGs with DNAm aging were proximate to GRHL2 5’UTR (z = 12.5), ADRB1 exon (z = 12.3), and PURA 3’UTR (z = −11.4) (**Figure 4A**). Aging-associated CpGs in deer blood were distributed in all genic and intergenic regions that can be defined relative to transcriptional start sites (**Figure 4B**). However, promoter regions had a higher proportion of hypermethylated CpGs compared to others. This result paralleled a higher positive association of CpG islands with age than non-island CpGs (**Figure 4C**).

**Figure 4:**
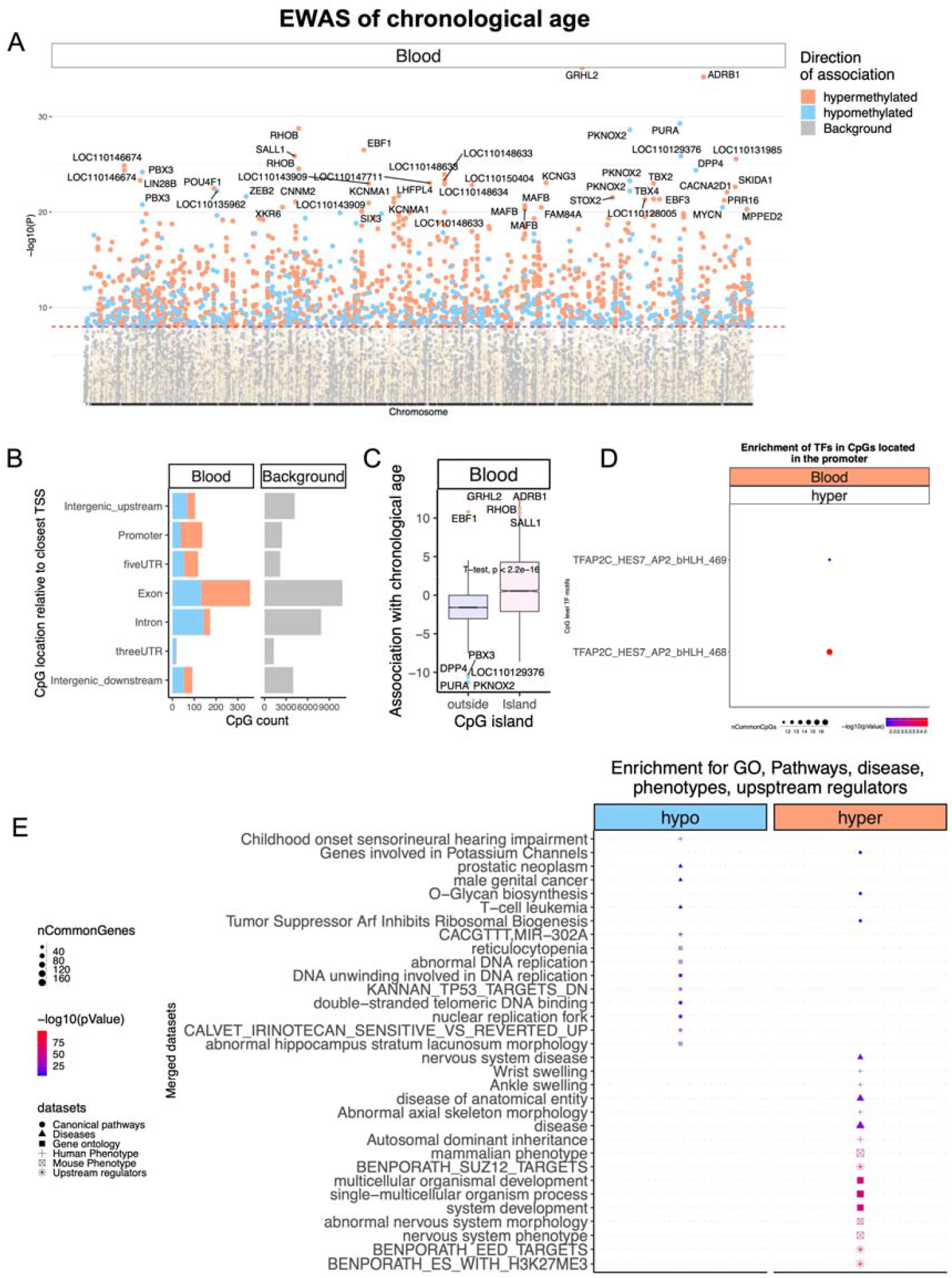
Epigenome-wide association (EWAS) of chronological age in the blood of roe deer. A) Manhattan plot of the EWAS of chronological age. Since the genome assembly is not available for roe deer, the coordinates are estimated based on the alignment of Mammalian array probes to White-tailed deer (Ovir.te_1.0) genome assembly, a related species to roe deer. The direction of associations with p < 10-8 (red dotted line) is highlighted by red (hypermethylated) and blue (hypomethylated) colors. The top 30 CpGs were labeled by the neighboring genes. B) Location of top CpGs in each tissue relative to the closest transcriptional start site. Top CpGs were selected at p < 10-8 and further filtering based on z score of association with chronological age for up to 500 in a positive or negative direction. The grey color in the last panel represents the location of 32767 mammalian BeadChip array probes mapped to Ovir.te_1.0 genome. C) CpG islands have a higher positive association with age (hypermethylation) than other sites. D) Transcriptional motif enrichment for the top CpGs in the promoter and 5’UTR of the neighboring genes. The motifs were predicted using the MEME motif discovery algorithm, and the enrichment was tested using a hypergeometric test. E) Enrichment analysis of the top CpGs in cat blood. The analysis was done using the genomic region of enrichment annotation tool (McLean et al. 2010). The gene-level enrichment was done using GREAT analysis (McLean et al. 2010) and human Hg19 background. The top 3 enriched datasets from each category (Canonical pathways, diseases, gene ontology, human and mouse phenotypes, and upstream regulators) were selected and further filtered for significance at p < 10-4.

Transcriptional factor enrichment analysis suggested TFAP2C motifs are hypermethylated with age in the leucocyte’s DNA of roe deer (**Figure 4D**). This motif is involved in cell-cycle arrest, germ cell development, and it is implicated in several types of cancer (Bryant et al. 2012; Penna et al. 2013). Understanding the functional outcome of this change in roe deer will require further studies.

Gene level enrichment analysis of the significant CpGs highlighted changes in development, the nervous system, O-Glycan metabolism, cancer, and immune system, all of which are associated with aging biology in humans and other species (**Figure 4E**). The analysis suggested that aging mediated hypermethylation is marked by H3K27Me3 and potentially regulated by polycomb protein EED targets. EED is a member of the multimeric Polycomb family protein complex that maintains the transcriptional repressive states of genes. These proteins also regulate H3K27Me3 marks, DNA damage, and senescence states of the cells during aging (Ito et al. 2018).

### CpGs whose aging patterns depend on sex

The epigenetic age acceleration was faster in male than in female roe deer. At genome-wide significance (p<10^−8^), age and sex altered DNAm of 1726 and 1022 CpGs, respectively (**Figure 5A**). This suggests both age and sex have a large effect size on DNAm levels. For sex, differentially methylated CpGs (DMCs) were biased in specific scaffold chromosomes, which are expected to be the homologs of sex chromosomes in humans and other mammals. For the interaction of age and sex, only three CpG proximate to RAI2 5’UTR, FAM155B exon, and ZIC3 exon had p-values <10^−8^. At a 5% false discovery rate (FDR), a total of 22 CpGs showed a statistically significant interaction of age and sex in roe deer blood samples. This suggests sex influences DNAm aging in roe deer.

**Figure 5:**
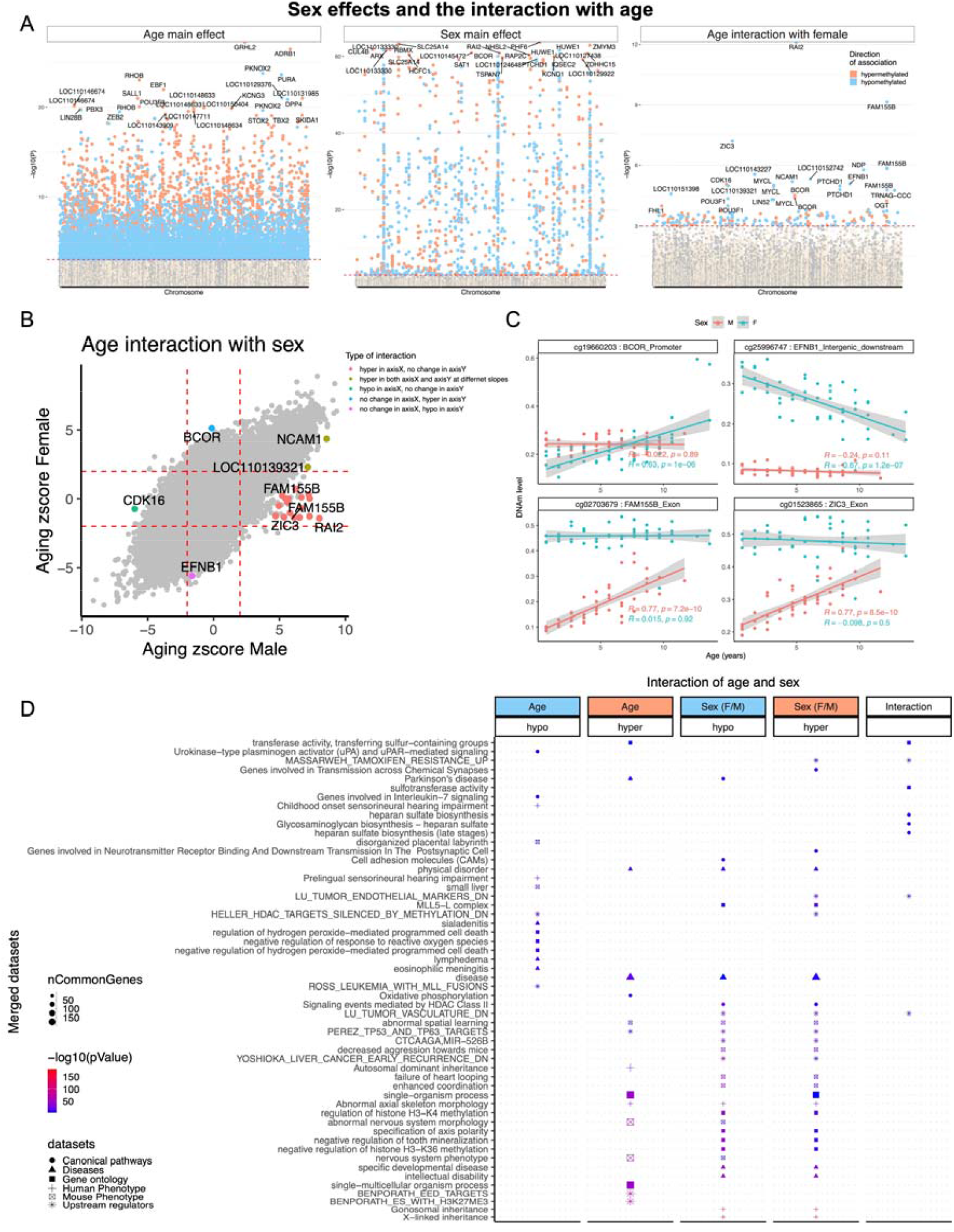
Sex influence on DNAm aging. A) Manhattan plots of DNAm aging loci that are shared between sexes (Aging main effect), basal sex differences (Sex main effect), and the interaction of sex and aging. The analysis is done by multivariate regression models with or without (to estimate the main effect) interaction term for age and sex. For sex, the male is the reference variable to estimate the direction of change. Sample sizes: Males, 45; Females, 49. The coordinates are estimated based on the alignment of Mammalian array probes to White tailed deer (Ovir.te_1.0) genome assembly. The red line in the Manhattan plot indicates p <1e-3. B) Scatter plots DNAm aging between male and female roe deer. The highlighted CpGs are the loci with statistically significant interaction between species at a 5% FDR rate. In total, five categories of interaction were defined based on the aging z-score of each sex. C) DNAm aging in selected loci with a statistically significant sex interaction. D) Enrichment analysis of the genes proximate to CpGs related to age (shared between sexes), sex, and age:sex interaction. The gene-level enrichment was done using GREAT analysis (McLean et al. 2010) and human Hg19 background. The top CpGs were selected at a 5% FDR rate and based on Beta values of association for up to 500 in a positive or negative direction.

We overlaid the CpGs that interacted between sexes on a scatter plot of aging z-scores of male and female roe deer. The analysis identified five kinds of interaction (Figure 5B). The strongest pattern was for CpGs that were hypermethylated with age in males, but not females. CpGs proximate to RAI2, FAM155B, and ZIC3 (**Figure 5C**) genes had the aforementioned pattern. In contrast, some CpGs such as BCOR promoter and EFNB1 downstream only influenced DNAm aging in females (**Figure 5C**).

## Discussion

Our findings highlight a very tight correlation between epigenetic and chronological age in two populations of roe deer intensively monitored in the wild. The quality of the fit of the selected model describing the relationship between epigenetic and chronologic age over the roe deer entire lifespan was particularly high (with a correlation coefficient of 0.975 leading to a R^2^ of 0.95; Median absolute difference: 0.588 years), which adds to the increasing evidence that the epigenetic tool constitutes an accurate method to estimate age in vertebrate populations in the wild (Paoli-Iseppi et al. 2017). So far, a wide range of technics based on tooth wear are generally used to assign age in wild mammals (Morris 1972, Pérez-Barbería et al. 2014 for a review in red deer, *Cervus elaphus*). In roe deer, tooth wear that leads the first molar height to decline with increasing age throughout the lifespan allows assessing age of individual roe deer (Veiberg et al. 2007). However, this method is much less accurate than the epigenetic clock (R^2^= 0.69 vs. 0.94 when a linear regression is used, Fig. S1).

Interestingly, the fit of the roe deer epigenetic clock outperforms the few epigenetic clocks previously developed from other mammalian populations. All studies performed so far have investigated the relationship between epigenetic and chronologic age through linear regressions, and did not account for non-linearities. The linear regression provided a better fit in roe deer here (i.e. a coefficient of correlation of 0.97) than that reported in humpback whales (*Megaptera novaeangliae*, correlation of 0.89, Polanowski et al. 2014); wood mice, (*Apodemus sylvaticus*, correlation of 0.92, Little et al. 2020) or Bechstein’s bat (*Myotis bechsteinii*, correlation of 0.80, Wright et al. 2018), which might be due to the broad age range we included in the analysis or to differences in the biological tissue used to extract DNA (e.g. leukocytes vs wing or ear punches). Overall, the epigenetic clocks used in our study constitute a particularly accurate method for estimating age in roe deer on the basis of leucocyte DNA.

We found that the relationship between epigenetic and chronologic ages was better described by a quadratic than linear model when juveniles were included, but any deviation from a linear model vanished when considering only adults. This discrepancy and the negative second order term of the quadratic model clearly indicate that the relationship between epigenetic and chronological ages is steeper in growing juveniles than in adults. A similar pattern has been reported in humans where the rate of change in DNA methylation profiles (also called ‘tick rate’) was higher during the developmental period than during adulthood, when a constant tick rate seems to be the rule (Horvath 2013). The growth period is associated with a high rate of mitotic division and constitutes a particularly demanding life stage in terms of resource allocation in mammals (Reiss 1989). Although overlooked for a while, the ageing consequences of a fast growth during early life are increasingly investigated (Metcalfe and Monaghan 2003) and recent evidence suggests that fast growth can shorten lifespan on the long-run (Lee et al. 2013; Kraus et al. 2013), even though the exact physiological mechanisms underlying this association are likely to be multiple and complex (Metcalfe and Monaghan 2003; Monaghan and Ozanne 2018). In roe deer, individuals reach their asymptotic mass around the age of 4 years in both males and females (Hewison et al. 2011). However, juveniles (i.e. at eight months when individuals are captured for the first time) have already gained about two-third of their adult body mass (Hewison et al. 2011), with a high amount of individual variation, which makes winter juvenile body mass a reliable measure of growth intensity. We found that the average age epigenetic acceleration increases with juvenile body mass, suggesting that individuals who allocate substantially in their growth are biologically older than their chronological age indicates, which might contribute to explain why, in roe deer, a fast-post-weaning growth is associated with a steeper rate of body mass senescence (Douhard et al. 2017). A positive association between DNA methylation profiles (as measured with the Horvath pan tissue clock) and height has also been observed among teenagers (Simpkin et al. 2016), which suggests that the discrepancies between biological and chronological age following fast growth might be widespread across mammals and also offer new perspectives for the study of the relationships between growth and ageing.

Despite the limited sample size, our analyses suggest that for a given age, male roe deer from the Chizé population are biologically older than females, a difference that is particularly pronounced at old ages. As the epigenetic age acceleration is associated with mortality risk in humans (e.g. Marioni et al. 2015), this result is in line with previous roe deer survival analyses, which showed that males at Chizé have a shorter lifespan than females - especially when born during years of strong environmental harshness (Garratt et al. 2015). Alike humans or laboratory rodents, there is compelling evidence that mammalian males in the wild display shorter lives than females (Lemaître et al. 2020c), as predicted by several (and non-mutually exclusive) evolutionary theories (e.g. heterogametic sex hypothesis, mother’s curse hypothesis, sex differences in life history strategies, see Austad and Fischer 2016; Marais et al. 2018 for reviews). However, the magnitude of sex differences in lifespan remains highly variable among populations and species (Lemaître et al. 2020c). This discrepancy might partly result from the high variation in environmental conditions faced by populations in the wild (Lemaître et al. 2020c; Tidière et al. 2020). More specifically, harsh environmental conditions (e.g. low availability in resources, high pathogen richness) are expected to amplify the survival cost of male reproductive expenditure (e.g. allocation to sexual traits, territory defence) due to the acute resource-based allocation trade-offs between reproduction and survival insurance mechanisms (Kirkwood and Rose 1991; Kirkwood 2017). At Chizé, environmental conditions are much harsher than at Trois Fontaines due to low quality resources, which might explain the clear epigenetic age acceleration in old males from this population.

More generally, our findings suggest that the epigenetic age acceleration might constitute a relevant biological marker of sex differences in health and biological conditions in mammals. For sex-related CpGs, the top enriched datasets were related to X-linked and Gonosomal inheritance. Other basal sex differences were related to the nervous system (e.g. synapse function), cognition (e.g. intellectual disability, spatial learning), and development (e.g. nervous system, teeth, muscles). For interaction, the enrichment analysis suggested a difference in heparan sulfate glucosaminoglycan biosynthesis between male and female aging. In humans, Alzheimer’s disease patients have higher distribution and localization of heparan sulfate glucosaminoglycan in neurons, microglia, and often colocalize with amyloid plaques (Su et al. 1992). Moreover, these macromolecules are involved in aging through neurogenesis (Yamada et al. 2017),and skin homeostasis (Bucay et al. 2020). Our results suggest a sex difference in heparan sulfate glucosaminoglycan biosynthesis during aging, which could also contribute to sex differences in human neurodegenerative disorders. Moreover, our analysis suggests DNAm underlie some of these differences. While mammalian females undeniably live longer than males (Lemaître et al. 2020c), studies that have sought to decipher the genetic and physiological correlates of these sex differences in survival have remained rather inconclusive. For instance, other biological markers of aging such as telomere length or immune performance do not show clear differences between males and females in wild mammals (Peters et al. 2019; Remot et al. 2020). We emphasize that using the epigenetic age acceleration across longitudinal follow-ups of mammalian populations does constitute to date the most promising approach to i) estimate accurately the chronological age of individual mammals, (ii) assess precisely sex differences in physiological condition, iii) disentangle the complex ecological and biological origins of these differences, and iv) establish reliable predictions in terms of individual trajectories. In the current context of a growing age in human populations associated with pronounced sex differences in the occurrence of age-associated diseases in the elderly (Austad and Fischer 2016; Clocchiatti et al. 2016), this research avenue is extremely promising.

## Author contributions

SH and JFL conceived of the study and wrote the article. The remaining authors helped with the statistical analyses and data collection. All authors reviewed and edited the article.

## Ackowledgments

This work was supported by the Paul G. Allen Frontiers Group (SH). We thank all the OFB staff, in particular Gilles Capron, Stéphane Chabot and Claude Warnant and the field volunteers for organizing the roe deer captures.

## Ethics

The protocol of capture and blood sampling under the authority of the Office National de la Chasse et de la Faune Sauvage (ONCFS) was approved by the Director of Food, Agriculture and Forest (Prefectoral order 2009–14 from Paris). The land manager of both sites, the Office National des Forêts (ONF), permitted the study of the populations (Partnership Convention ONCFS-ONF dated 2005-12-23). All experiments were performed in accordance with guidelines and regulations of the Ethical Committee of Lyon 1 University (project DR2014-09, June 5, 2014).

## ELECTRONIC SUPPLEMENTARY MATERIAL

**Table S1:** Model selection procedure from the set of linear models fitted to test the relationship between DNAm age and chronological age for all individuals in the dataset (A) or for adults only (B). The selected model is highlighted in bold, k is the number of parameters in the model, ΔAIC is the difference in AIC between the candidate model and the selected model. The AIC weight (AICw) is calculated to measure the relative likelihood that a given model is the best among the set of fitted models.

| | | | k | AIC | $\Delta AIC$ | AICw |
| --- | --- | --- | --- | --- | --- | --- |
| **A** | All individuals<br>(N=90) | Constant | 2 | 448.09 | 260.46 | 0.00 |
|  |  | Linear | 3 | 200.72 | 13.09 | 0.00 |
|  |  | **Quadratic** | **4** | **187.63** | **0.00** | **1.00** |
| **B** | Individuals older<br>than 1-year-old<br>(N=82) | Constant | 2 | 334.37 | 165.86 | 0.00 |
|  |  | **Linear** | **3** | **169.06** | **0.55** | **0.43** |
|  |  | Quadratic | 4 | 168.51 | 0.00 | 0.57 |

**Table S2:**
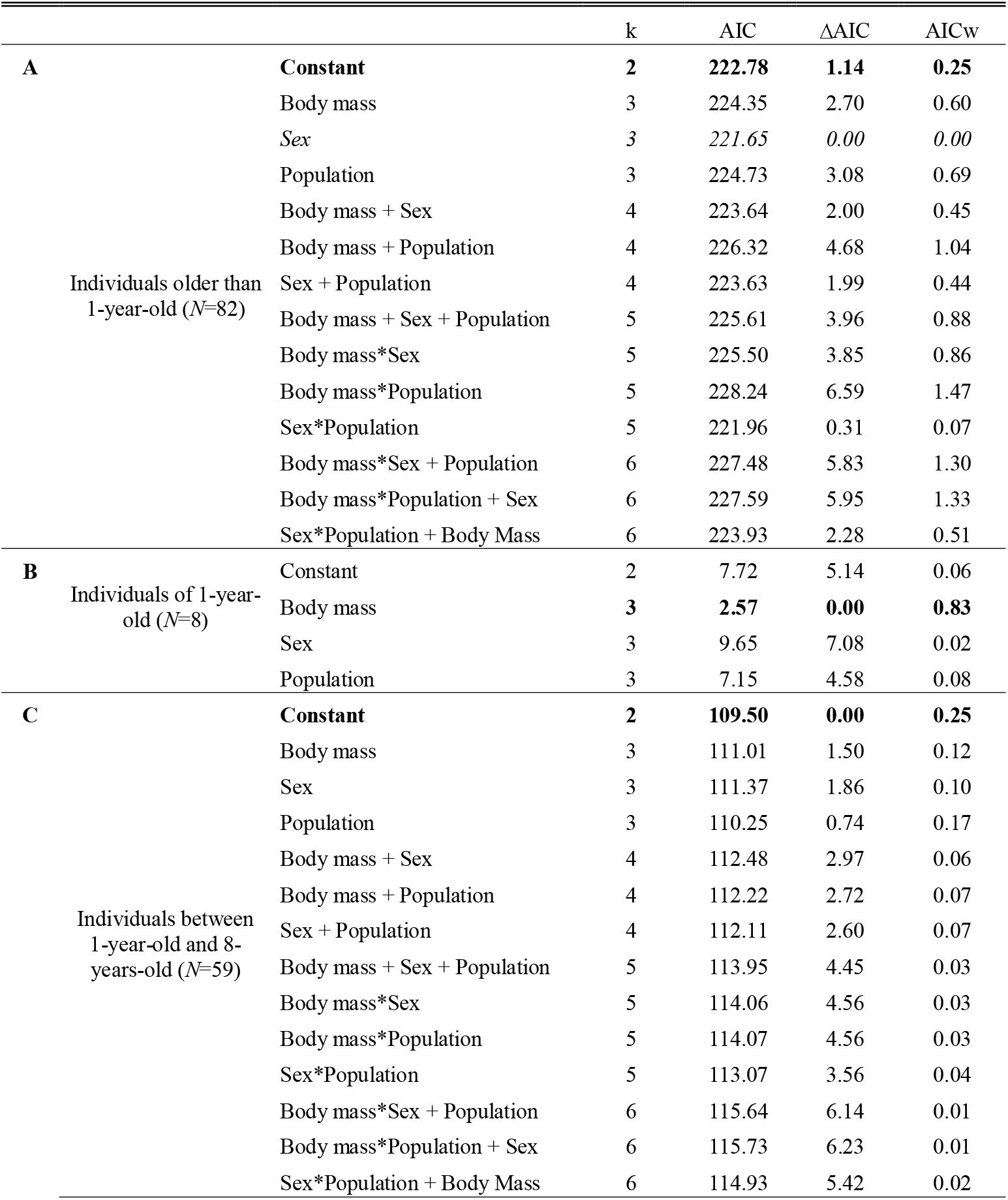
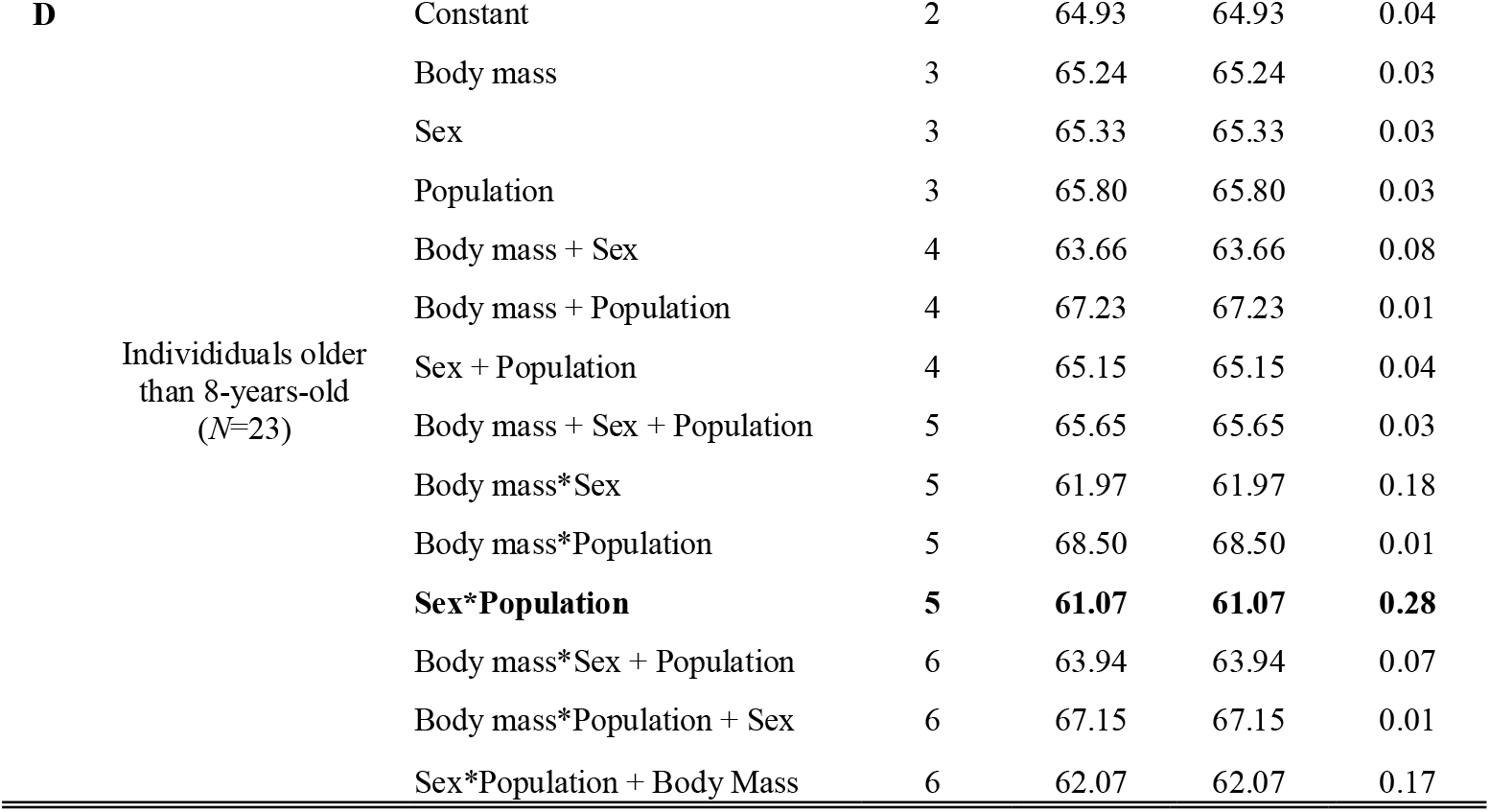
Model selection procedure from the set of linear models fitted to test the relationship between the Average age acceleration in the epigenetic clock and chronological age, sex, population and body mass for adults (A), juveniles (B), Prime-aged individuals (C) and senescent individuals (D). The selected model is highlighted in bold, k is the number of parameters in the model, ΔAIC is the difference in AIC between the candidate model and the selected model. The AIC weight (AICw) is calculated to measure the relative likelihood that a given model is the best among the set of fitted models.

**Table S3:** Quadratic model describing the relationship between DNAm age and chronological age in roe deer (*N*=90). Contrary to Table 1, the model was fitted with roe deer identity included as a random effect. Results are qualitatively unchanged (\**p*<0.05, \*\**p*<0.01,*** *p*< 0.001).

|  | Estimate | SE | t |
| --- | --- | --- | --- |
| Intercept | 0.28 | 1.31 | 1.39 |
| Age | 1.13 | 0.08 | 14.90*** |
| Age<sup>2</sup> | -0.02 | 0.006 | -4.02*** |

**Table S4:** Parameters of the models including the interaction between chronological age and sex using the female (a) or the male (b) clock; and of the models including the interaction between chronological age and population using the Trois Fontaines (c) or the Chizé (d) clock. Contrary to Table 2, these models were fitted with the roe deer identity as a random effect. Results are qualitatively unchanged (\**p*<0.05, \*\**p*<0.01,*** *p*< 0.001).

| Dependent Variables |  | Estimate | SE | t |
| --- | --- | --- | --- | --- |
| (a) DNAm age [Female epigenetic clock] | Intercept | 1.03 | 0.2 | 5.09*** |
|  | Age | 0.85 | 0.03 | 30.91*** |
|  | Sex | 0.7 | 0.3 | 2.41** |
|  | Age * Sex | -0.07 | 0.04 | -1.57 |
| (b) DNAm age [Male epigenetic clock] | Intercept | 3.32 | 0.18 | 18.40*** |
|  | Age | 0.48 | 0.02 | 19.86*** |
|  | Sex | -1.92 | 0.27 | -7.10*** |
|  | Age * Sex | 0.25 | 0.04 | 6.06*** |

**Figure S1:**
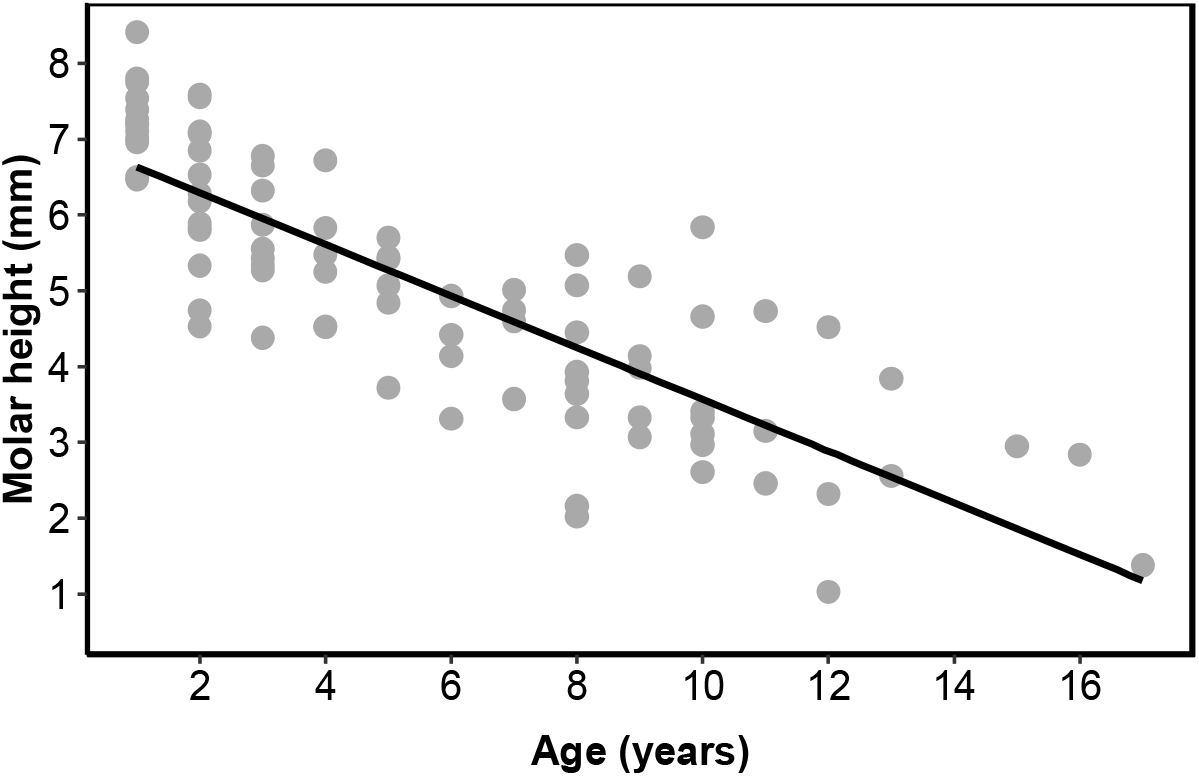
Relationship between molar height (M1, in mm) and age (in years) for roe deer from Chizé and Trois-Fontaines (slope ± se: −0.34 ± 0.02, *N*=88, R^2^= 0.69).

